# The isolated *Stachel* peptide of the adhesion G protein-coupled receptor GPR126 is intrinsically disordered

**DOI:** 10.64898/2026.01.16.700034

**Authors:** Tucker Shriver, Sandra Berndt, Scott A. Robson, Austin D. Dixon, Ines Liebscher, Joshua J. Ziarek

**Affiliations:** Department of Pharmacology, Northwestern University Feinberg School of Medicine, Chicago, Illinois, USA; Rudolf Schönheimer Institute of Biochemistry, Medical Faculty, Leipzig University, Leipzig, Saxony, Germany; Biologics Discovery Sciences – Biologics Production, AbbVie, Chicago, Illinois, USA

**Keywords:** adhesion GPCRs, ADGRG6, peptide NMR, structure, ligand, disordered, disorder-to-order transition, order-to-disorder transition

## Abstract

Several members of the adhesion subfamily of G protein-coupled receptors (aGPCRs) are capable of self-activation by an internal agonist sequence (aka the *Stachel*) that’s exposed upon removal or conformational changes of the N-terminal fragment of the receptor. Synthetic peptides derived from the *Stachel* sequence can be used as exogenous agonists. In the inactive form, the *Stachel* is sequestered as the β13-strand within the GPCR Autoproteolysis-INducing (GAIN) domain, but it engages the seven transmembrane region as a helix when it is either an intramolecular sequence or a synthetic peptide. Little is known about the molecular details underlying this transition, but we hypothesize that a disordered conformation is central to this intermediate state in receptor activation. Despite the primarily helical *Stachel* AlphaFold3 and Pepfold4 models predicted with high confidence for the entire aGPCR subfamily, disorder predictions and biophysical experiments reveal a predominantly disordered conformation in solution. Investigating the ADGRG6/GPR126 *Stachel* peptide, circular dichroism (CD) and nuclear magnetic resonance (NMR) experiments reveal a predominantly random coil conformation in aqueous buffer, polar detergent micelles, and zwitterionic lipids. Titration of trifluoroethanol uncovered a two-state equilibrium between an unfolded and helix-containing conformation with NMR localizing a single-turn helix to residues L846-L849. Taken together, these data indicate the ADGRG6/GPR126 *Stachel* peptide is primarily disordered, but small populations may adopt a helix-containing conformer that seems to support a conformational-selection activation mechanism.

## INTRODUCTION

The thirty-three member aGPCR subfamily^1^, first characterized in 1995, represents the second largest subclass of GPCRs^2^, and yet their mechanism of activation remains arguably the most understudied^3,4^. N-terminal to the superfamily defining seven transmembrane (TM) domain, most adhesion GPCRs possesses a variable set of domains (e.g. CUB, PTX, SEA, *etc.*^5^) that span hundreds to thousands of residues in length (Fig. 1A). All aGPCRs except one encode a GPCR autoproteolysis-inducing (GAIN^6,7^) domain directly adjacent to the TM. This GAIN domain mediates self-cleavage during biogenesis for most but not all aGPCRs to result in two distinct polypeptide fragments being expressed at the cell membrane^8,9^. These two fragments, the N-terminal fragment (NTF) and the C-terminal fragment (CTF), remain non-covalently associated ^10^. The NTF comprises the GAIN domain up to the conserved cleavage motif and any other N-terminal domains whereas the CTF contains the intramolecular *Stachel* agonist sequence and the TM domain. One mechanism of receptor activation is initiated when the NTF binds cellular factors that confer mechanical force upon the NTF, which result in NTF release or conformational changes, exposing the intramolecular agonist *Stachel* peptide ^11–13^ (Fig. 1B). The newly exposed, now N-terminal *Stachel* peptide is then unrestricted to self-associate with the 7TM orthosteric pocket to promote activation. Based on structures of several GAIN domains^14^, a putative activation mechanism for the *Stachel* sequence would require conversion from a *β*-stranded form when bound to the GAIN domain to a mixture of helical and random coil secondary structures when bound to the receptor. While these detail the specific interactions of the *Stachel* peptide in its two primary molecular environments, they reveal little about the underlying transition between them. When considered outside the context of the GAIN domain, the ADGRG6/GPR126 *Stachel* peptide sequence has been previously^15^ predicted to adopt a secondary element-turn-secondary element architecture, the presence of which would influence the resulting mode of activation (induced fit vs conformational selection).

**Figure 1.**
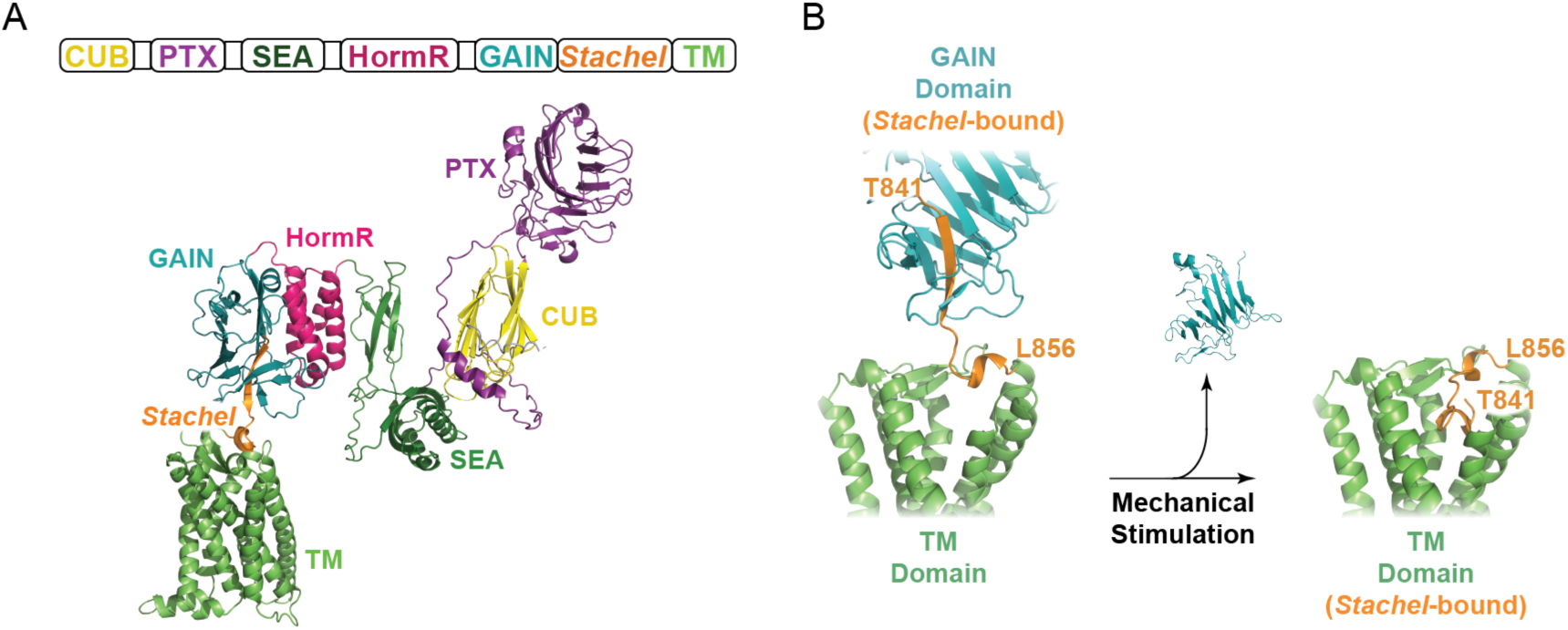
Structural models of hGPR126 cryptic and exposed *Stachel* peptide. A) Alphafold3 model of wild-type hGPR126 with each N-terminal domain uniquely colored to match the primary sequence. The composite model presented here is an overlay of AF3 models of the NTF-Stachel and GAIN-CTF (sequences in Table S2) aligned by the GAIN domain. B) During the process of activation, it is hypothesized that the *Stachel* peptide (orange) transitions from its position as the 13^th^ *β*-strand of the GAIN domain (left) to a site buried in the TM domain (right), though no structure of the TM-bound GPR126 receptor has yet been obtained. The goal of this study is to characterize the conformation(s) between these two states.

The actual secondary structure of this peptide has not been previously determined in the solution state. Here, we modeled all thirty-three human aGPCR *Stachel* peptides, in isolation, using Pepfold4. We determined the stachel sequences based on gpcrdp.org numbering and determined that the majority adopted an ordered state containing a full or partial helix. We next assessed the degree of disorder predicted with a focus on GPR126. Next, circular dichroism (CD) measurements of the isolated GPR126 *Stachel* peptide were collected in aqueous buffer, polar and zwitterionic membrane mimetics, and the secondary structure-inducing cosolvent trifluoroethanol (TFE). Spectral deconvolution was used to determine the percent composition of each secondary structure; TFE titration illustrated a two-state equilibrium between a disordered and a single-helix conformer. Natural abundance nuclear magnetic resonance (NMR) was then used to assess the helix location. These comparisons revealed a clear induction of a single turn helix within L846-L849, likely acting as a key conformation determinant for the entire peptide and enabling binding to the transmembrane (TM) domain.

## RESULTS

### AlphaFold3 and Pepfold4 Model a Structured *Stachel* Peptide Despite Disorder Prediction

We used AlphaFold3 (AF3)^16^ to model the GAIN domain-bound (i.e. inactive) and receptor-bound (i.e. active-like) GPR126 *Stachel* peptide, which revealed a dramatic secondary structural change from *β*-strand to *α*-helix (Fig. 2A). This change in secondary structure is predicted to be shared among all cleavable aGPCRs^17–19^. With the goal of structurally-characterizing the intermediate state ^15^, we first measured the conservation scores for all 33 human aGPCR *Stachel* peptides with adjacent stalk region. These sequences are shown in Table S1 and were defined as the residue C-terminal to the proteolytic cleavage site (or the beginning of the sequence for ADGRA1) through the first residue of TM1, as defined by GPCRdb.org. Not all aGPCRs need their *Stachel* sequences for activation and thus some sequences have diverged sufficiently from those needed for activation. aGPCR phylogeny has been based on conservation of the transmembrane domain, not the *Stachel* sequences so to account for the variance, conservation scores (CS) were calculated and transformed to percent divergence (1-CS) and all sequences (6 total) with divergence scores greater than the median plus median average deviation were not modeled. The subsequent 27 sequences were then used in Pepfold4 to compare the predicted secondary structures. As shown in Figure 2.B, 7 sequences were predicted to adopt a *β*-stranded conformation, 6 a partial or broken helix, and the majority (14) a full helix. We then calculated the consensus *Stachel* sequence as TSFAVLMxLS where x is unspecified. Only top scoring pepfold4 models were considered, but analysis of other top 5 models shows some population of both helical and stranded conformations, suggesting the ability of many to transition between them. We also considered the effect of varying sequence length on structural predictions for ADGRG6 *Stachel* sequence and found no variation in secondary structure as a function of sequence length (data not shown). This aligns with previously published data which shows a mostly linear decrease in GPR126 activation (cAMP accumulation) as a function of sequences length, with shorter sequences showing reduced activation^20^.

**Figure 2.**
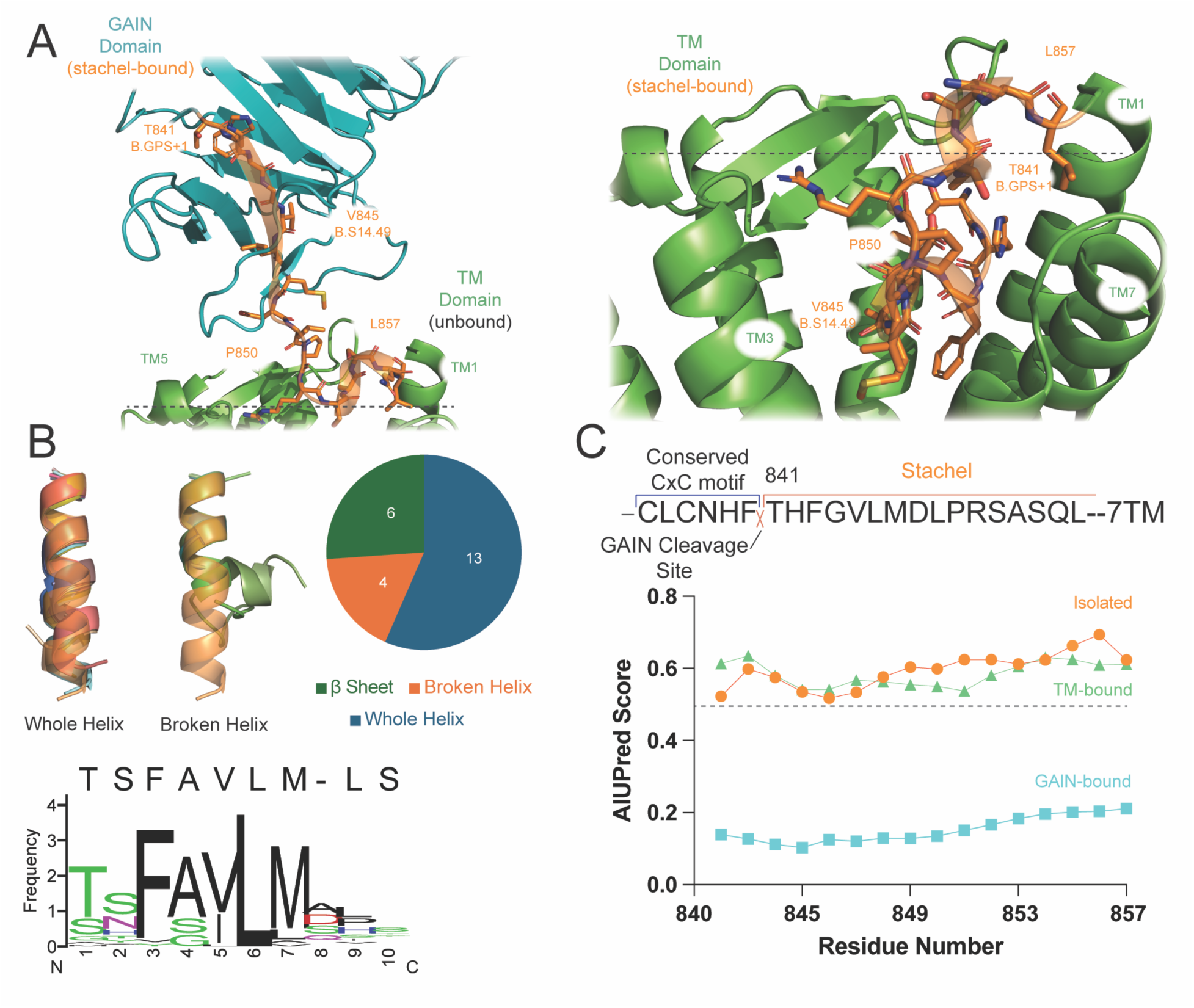
Models suggest a structured *Stachel* despite predicted disorder. A) Key contacts between the Alphafold3-generated hGPR126 *Stachel* peptide (T841-L856; orange) when modeled with the GAIN domain (cyan; left) and the TM (green; right). Residues labeled according to GPCRdb^22^. B) Pepfold4 models of the 27 human aGPCRs *Stachel* peptides found to have appropriate conservation scores (Fig. S1). Overlays show the ADGRG6 *Stachel* in orange with 60% opacity compared with whole and broken helix groups in 100% opacity. Consensus sequence shown below generated from these same 27 sequences. C) AIUPred disorder scores for *Stachel* region (T841-L856) in the inactive (GAIN-bound; residues D665-L856; cyan), intermediate (isolated; residues T841-L856; orange), and active-like (TM-bound; residues T841-C1221; green). Only residues of the *Stachel* region are reproduced although the entire indicated sequence was included in the prediction (Table S2). Scores > 0.5 qualify indicate disorder.

We hypothesized that the *Stachel* peptide’s secondary structure was modulated by neighboring domains with which it interacts. We used the AIUPred server^21^, a highly accurate deep neural network for disorder prediction, to assess the GPR126 *Stachel* peptide in three states (Fig 2C, SI Table 2): inactive (GAIN-bound), intermediate (isolated), and active-like (TM-bound). Contrary to the AF3 prediction (Fig. 2B), the isolated *Stachel* peptide shows almost an entirely disordered character (Fig. 2C). Whereas inclusion of the N-terminal GAIN domain dramatically orders the Stachel as expected based upon the published structures^14^, concatenation with the TM domain has little-to-no impact on predicted disorder (Fig. 2C). This would suggest there is some dynamic feature of the *Stachel* sequence that is not captured by crystal structures or modeling.

### GPR126 *Stachel* Peptide Remains Disordered in the Presence of Membrane Mimetics

Far-UV circular dichroism (CD) was used to determine the average secondary structure of the GPR126 *Stachel* peptide under varying solution conditions^23,24^. CD measures the differential absorption of left- and right-circularly polarized light, reported as ellipticity, across a range of wavelengths. The regular arrangement of chromophores contained in secondary structural elements absorb this light with recognizable spectral signatures. The resulting CD spectra are population-weighted, which makes deconvolution necessary when a target protein contains multiple secondary structure elements. Numerous reference libraries of proteins with known structure have been collated to emphasize a particular property (e.g. MW, integral membrane protein, etc). The beta structure selection (BeStSel)^25^ webserver was used for its purported ability to more accurately detect *β*-stranded character, a noted difficulty with other algorithms. First, a spectrum collected in sodium phosphate buffer was predicted as 44% stranded, 14% turn, and 42% other; Figure 3A shows the experimental CD trace along with the reconstructed trace derived from BeStSel deconvolution. Next, we tested the effect of environment on GPR126 structure. To mimic the lipid bilayer, we collected spectra using nonionic (lauryl maltose neopentyl glycol; LMNG) detergent and zwitterionic (1,2-dihyexanoyl-sn-glycero-3-phsophocholine; DHPC) lipidic micelles. Both membrane mimetics led to a subtle (5-8%) reduction in β-stranded structure that was accompanied by an increase in random coil character (Fig. 3B). This would suggest that increased lipophilicity has a minimal effect on reordering the *Stachel* secondary structure and that hydrophobic effect is unlikely to mediate any structural rearrangement.

**Figure 3.**
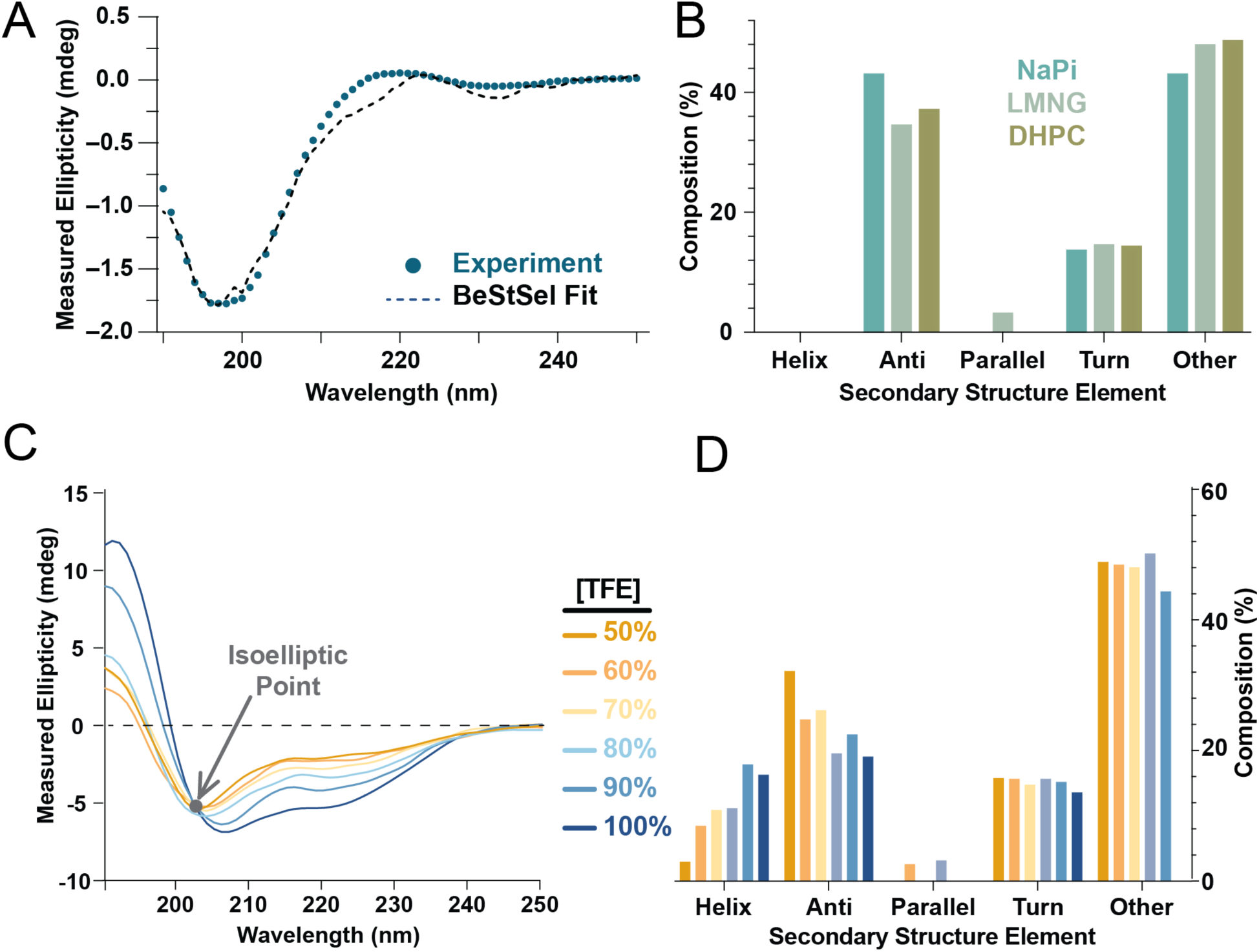
The effect of lipids, detergents and cosolvents on GPR126 secondary structure. A) Representative far-UV CD spectrum of hGPR126 *Stachel* peptide in sodium phosphate overlaid with BeStSel-reconstruction (solid line). B) Comparison of BeStSel deconvolution profiles for NaPi, LMNG and DHPC. C) Spectral overlay of TFE concentration series. The spectra overlap reveals a isoelliptic point at ∼203 nm, which indicates a two-state equilibrium. D) Deconvolution of TFE concentration series suggests the co-solvent induces helicity within ∼20% (3.2 AA) of the GPR126 *Stachel* peptide.

### Trifluoroethanol Reversibly Induces Local *Stachel* Helicity

2,2,2-Trifluoroethanol has a long history as a co-solvent in peptide studies^26^ that stabilizes secondary structure formation^27^. We collected CD spectra of GPR126 *Stachel* peptide in the presence of increasing TFE concentrations from 50-100% (Fig. 3C). Spectral overlays reveal an isoeliptical point at ∼202nm, indicative of a two-state conformational equilibrium without any significant intermediate species^28^. The BeStSel deconvolutions reveal that TFE promotes a transition from ∼20 % β-stranded to 20% helical character (Fig. 3D). As there are no appreciable change in other structural elements, we hypothesize this reflects the architectural transformation within a single, approximately four residue region of the GPR126 peptide reminiscent of receptor-bound *Stachel* peptide structures (7SF8^29^, 7WUJ^30^, 7EPT^10,31^).

### NMR Defines Region of TFE-Induced Helicity

To determine which GPR126 residues adopt a helical conformer, we employed nuclear magnetic resonance (NMR). Samples were prepared in both i) NaPi (pH 7.4) and ii) 90% TFE in NaOAc (pH 4.5). Resonances were assigned to 85% completion for both conditions using a suite of standard 2D homo- and heteronuclear correlation experiments (Fig. 4A). Following resonance assignment, we measured the secondary chemical shifts^32^, temperature coefficients^33^ and NOE-contacts^34^ to assess secondary structure formation.

**Figure 4.**
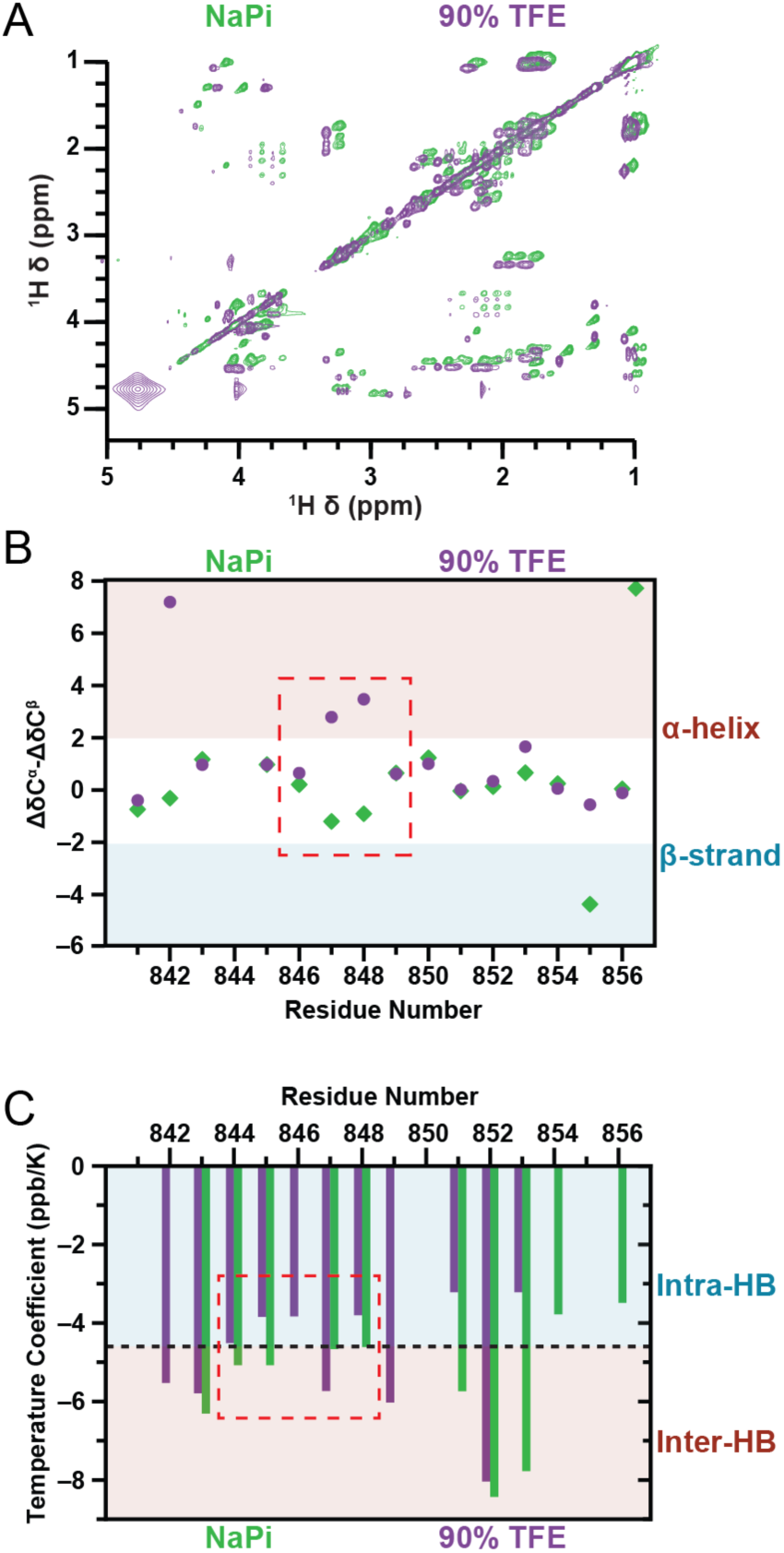
NMR Secondary chemical shifts and temperature coefficients refine the helix location. A) Representative TOCSY spectra of hGPR126 *Stachel* in 20 mM NaPi (pH 7.4) in green and 90% TFE in 20 mM NaOAc (pH 4.5) in purple. B) Secondary shift changes indicate an increased helical propensity at residues M847 and D848 in 90% TFE (purple dots) compared to NaPi (green diamonds). C) Amide temperature shift coefficients show a slight induction of helicity within residues L846-L849 as well (purple; TFE, green; sodium phosphate). No resonances were observable for T841, P850 or Q855.

Digital repositories of chemical shift information (BMRB^35^) have enabled informatics-based characterization of various protein NMR quantities like average chemical shift. These averaged, random coil values act as a reference for other experimentally determined shifts for each residue and atom type. The chemical shift value reflects how the local electronic environment (i.e. electron density) around a nucleus (de)shields it from the external magnetic field. Each atom of an amino acid has a characteristic “random coil” chemical shift that reflects its unstructured, flexible and unfolded state. As part of a polypeptide the backbone torsion angles, hydrogen bonding, ring currents, and other local environmental effects further influence the electronic environment as so called “secondary chemical shifts” ^36–38^. The secondary chemical shift is rigorously defined as the change in chemical shift (Δδ) between the observed and random coil chemical shifts. For example, the C^α^ atoms in α-helices tend to have positive secondary chemical shifts while those in β-strands have negative secondary chemical shifts; as C^β^ atoms show exactly the opposite behavior, taking the difference between ΔδC^α^ and ΔδC^β^ secondary chemical shifts is a common way to assess local secondary structure. Using this method, there is a subtle induction of helical character within residues L846- L849 (Fig 4B) as evidenced by the increase in average secondary chemical shift for the TFE condition.

Another parameter that qualitatively correlates with helix formation is the amide proton temperature coefficient (ΔδH^N^/ΔT). Amide protons rapidly exchange with water and thus the magnitudes of change in their temperature-dependent chemical shift are indicative of hydrogen bond formation within protein secondary structures^39^. NMR studies on a set of folded proteins, which also had high resolution crystal structures, demonstrated that more than 85% of the time these structured amide protons had temperature coefficients > −4.6 ppb/K ^39^. Stachel residues 844-846 and 848 possess temperature coefficients > −4.6 ppb/K consistent with intramolecular hydrogen bonds and formation of a single helix (Fig 4C).

Finally, we performed nuclear Overhauser effect (NOE) spectroscopy (NOESY) to provide data on internuclear distances. NOEs are a consequence of dipole-dipole coupling between different nuclear spins where the peak intensity is related to internuclear distance; in practice, NOEs are only observable for nuclei within ∼ 6 Å distance. Each of the various secondary structural elements produce distinct short-range ^1^H-^1^H NOE patterns^40^. The *Stachel* peptide shows a number of distant, through-space contacts (853A-HN to 844G-HN, 856L-HN to 845V-HN, Fig. 5A left panel) consistent with a disordered structure flexible enough to pivot around P850 and enable proximity of the N- and C-termini. In TFE, the *Stachel* peptide clearly produces NOEs consistent with the formation of a helix from approximately L846-L849 (Fig. 5A, right panel). It should be noted that all NOEs are population-weighted by the underlying conformers (e.g. folded or unfolded) and that some expected NOEs are intrinsically weak; taken together, it’s unlikely for a complete NOE set to be observed over the whole length of the helix^40^. Figure 5B details the complete summary of secondary structure data for the *Stachel* peptide in 90% TFE, showing numerous (i,i+1) amide-to-amide NOEs as well as singular (i,i+2), (i,i+3) and (i,i+4) amide-to-amide NOEs and multiple H_!_-to-amide NOES. In addition, temperature shift and secondary chemical shift data are also shown, further highlighting the formation of a helical element with residues L846 to L849.

**Figure 5.**
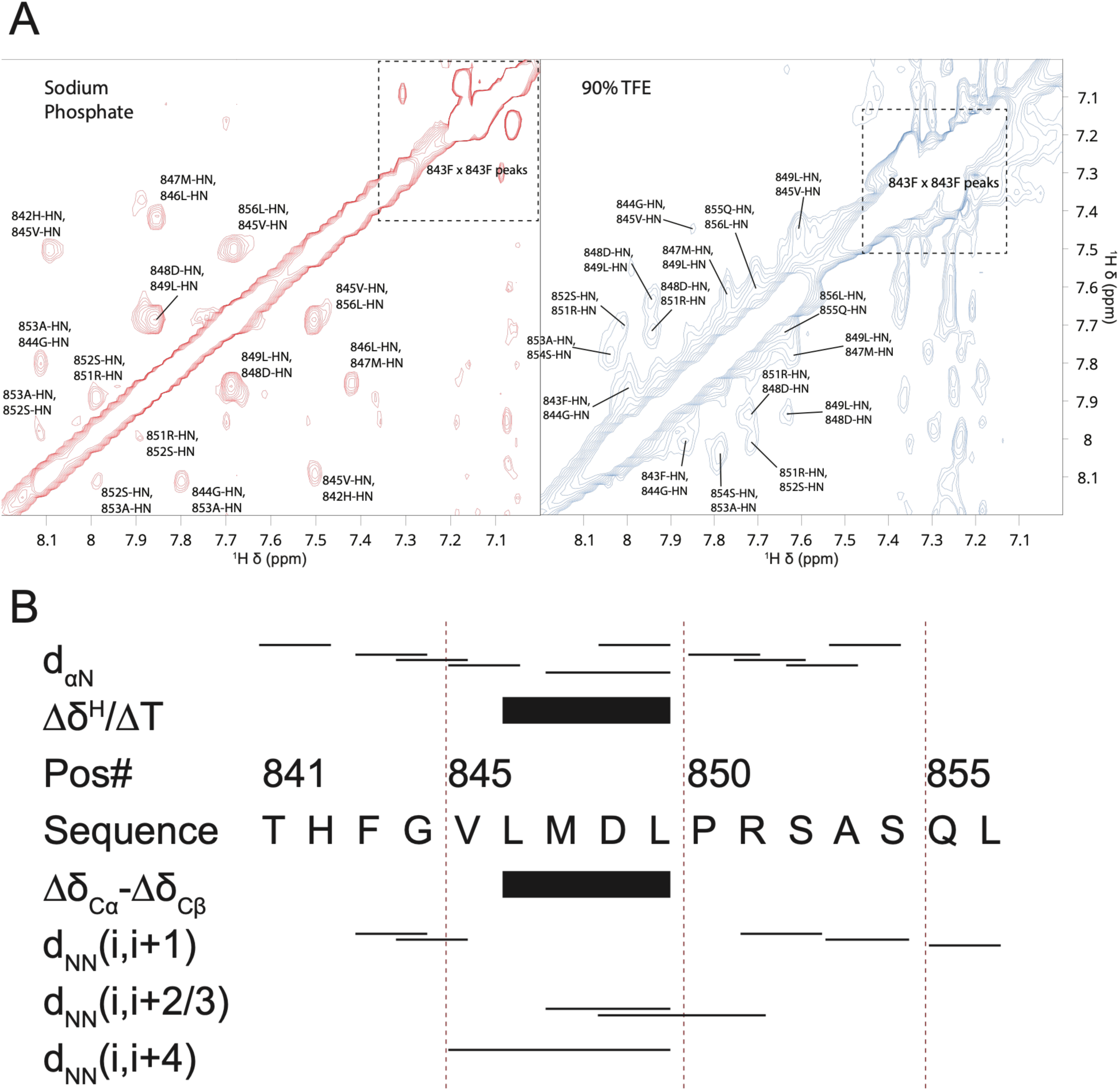
Summary of hGPR126 *Stachel* peptide NMR observables consistent with helix formation in TFE. A) Amide-amide NOESY contacts for sodium phosphate (left, red) and 90% TFE (right, blue). Thin lines represent NOE contacts between the residues at the left and right sides of the line. Black boxes summarize temperature coefficient (ΔδH^N^/ΔT) and secondary chemical shift (ΔδC^α^ and ΔδC^β^) values that are consistent with helix formation.

## DISCUSSION

We have shown that despite high AIUPred-predicted disorder scores for many aGPCR *Stachel* peptides, Pepfold4 predicted that all adopt structured conformations with relatively high confidence. GPR126, in particular, is predicted by Pepfold4 to adopt a helix from residues L846 to L849. To probe this seeming contradiction, we examined the effects of exposure to both polar (LMNG) and zwitterionic (DPC) membrane mimetics on its global secondary structure using CD and found little to no effect. We did observe a reversable induction of helicity when the GPR126 *Stachel* peptide exposed to increasing concentrations of known secondary structure induction reagent TFE, as measured by CD. The mean residue molar ellipticity indicates ∼20% helix formation, or approximately 3-4 amino acids, likely corresponding to a single helical turn. NMR analysis revealed, through both a reduction in amide protein exchange (temperature coefficient) and an increase in backbone secondary chemical shift changes, that the helical segment likely forms between L846-L849. Further work is needed to determine if these TFE-induced secondary structure changes represent physiologically relevant data, and future studies will include the determination of inter- and intrapeptide contacts within the TM-bound, *Stachel* peptide complex via NMR-based techniques like WaterLOGSY and transfer NOEs.

While our studies may not be indicative of the *specific* mechanism for GPR126, we believe they are indicative of a general mechanism. The presence of multiple bound-state structures for various peptides and their cognate receptors^41–44^ highlight the inherent dynamics required for some peptide-receptor pairs. Kinetic on- and off-rates are deeply tied to the modulations of accessible peptide conformational space. Interactions with both the membrane and membrane-bound receptors change the peptide’s propensities for different secondary structures. This can be seen in the emergence of regions which seed bound-state structures like in dynorphin^45^ and ghrelin^46^ binding. In both cases, a hydrophobic core at the interior of the peptide sequence is the first region to develop helical character on path to the bound state and lower the activation barrier for complete binding. Even when bound, dynorphin shows considerable dynamics at its N-term, going against traditional ideas of message-address binding for opioid peptide ligands^47^. Recently published work^48^ on ADGRD1 molecular dynamics simulations has shown the presence of multiple, low-energy structural intermediates on pathway to binding of the *Stachel* sequence to a G_s_-bound receptor complex. These intermediate states showed varying levels of secondary structure formation induced by interactions with neighboring residues on the extracellular interface.

This expanded understanding of the role of dynamics plays within receptor binding seems to further play out for aGPCRs. Based on our data, we posit the *Stachel* peptide undergoes an order-to-disorder transition upon GAIN domain release followed by a disorder-to-order transition during TM binding and receptor activation. The formation of electrostatic contacts between solvent and the peptide backbone is likely potentiated by the helix forming unit of L846-L849, and the proline immediately after this unit (P850) likely acts as a hinge residue, enabling the intercalation of the peptide into the cytosolic binding cavity of the TM domain. This aligns with other data suggesting that the P850A^12^ mutation serves to activate GPR126 *Stachel* binding, likely through lowering interconformer energy barriers associated with proline isomerization.

## MATERIALS AND METHODS

### GPR126 *Stachel* Peptide Synthesis

Solid phase peptide synthesis was performed on an automated peptide synthesizer MultiPep from Intavis AG (Köln, Germany) using standard Fmoc-chemistry. The final side chain deprotection and cleavage from the solid support employed a mixture of TFA, water and thioanisole (95:2.5:2.5, Vol %) for the peptides. The peptides were purified to >95 % purity using preparative RP-HPLC (Shimadzu LC-8, Duisburg, Germany) equipped with a PLRP-S column (300 x 25 mm, Agilent, Waldbronn, Germany). For both analytical and preparative use, the mobile phases were water (A) and acetonitrile (B), respectively, each containing 0.1 % TFA. Samples were eluted with a linear gradient from 5 % B to 90 % B in 30 min for analytical runs and in 90 min for preparative runs. Finally, all peptides were characterized by analytical HPLC (Agilent 1100) and MALDI-MS (Bruker Microflex LT, Bremen, Germany), which gave the expected [M+H]^+^ mass peaks.

### AlphaFold3 and Pepfold4 Modeling

A first run of the full-length GPR126 structure posed the NTF in a non-physiologic, lipid-embedded conformer. Therefore, the full length GPR126 structure was obtained by using AF3 to separately model the GPR126-NTF domain and a GAIN-CTF construct with a membrane mimetic of fifty molecules each of oleic and palmitic acids. These two structures were then aligned by their shared GAIN domains. This structure is consistent with the orientation of the various domains in the published NTF structure from *D. rerio*^14^. Sequences used for submission are detailed in Table S2.

*Stachel* peptide sequences were determined based on GPCRdb.org numbering (Table S1), starting at the GPS+1 site (or N-terminus for ADGRA1) and continuing through the first residue of TM1. Sequences were aligned via Clustal Omega webserver^49^ and only those *Stachel* sequences with divergence scores (1 - conservation score%) less that one median standard deviation away from the median score were kept, total of 27 (Fig S1). The Pepfold4 webserver was used to model these remaining *Stachel* sequences over AF3 due to its more specific training for peptides. For each sequence, 100 models were generated using 30,000 Monte Carlo steps at a simulation temperature of 370 K in a pH 7.5 environment with 20 mM ionic strength and no Debye-Huckel contribution. Top scoring models were aligned in PyMOL^50^.

Disorder Prediction: The AIUPred server^21^ was used to assess disorder prediction through a pairwise potential energy-based algorithm. Calculations were carried out with default smoothing.

### Circular Dichroism

Samples were prepped at 60 uM peptide in PBS (pH 7.4) or 60 uM peptide in 150 mM NaCl, 20 mM sodium acetate (pH 4.5) for TFE samples. PBS was supplemented with 1.26 mM DHPC and 0.009 mM LMNG corresponding to 90% (w/v) their respective CMC. All samples were degassed for twenty minutes immediately prior to measurement. All measurements were carried out on a Jasco J-1715 circular dichroism spectropolarimeter with a Peltier temperature controller under external nitrogen flow using a 1cm pathlength quartz cuvette. All spectra were collected over a range of 190-260 nm. Each CD trace represents the average of twenty scans. Buffer background spectra were subtracted from experimental spectra before analysis. The BeStSel server^25^ was used for spectral deconvolution and secondary structure estimation.

### Nuclear Magnetic Resonance Assignment

Assignment samples were prepared at 2 mM in PBS with 5% D_2_O and 20 uM DSS. All spectra were acquired at the MetaCyt Biomolecular NMR Laboratory at Indiana University on a 600MHz Bruker Avance NeoII console with a triple resonance (HCN) cryoprobe. Spectra were processed using nmrPipe^51^ and assigned using CCPNMR^52^, both services hosted by nmrBox^53^. The peptide was also assigned using the same workflow in a 90% (v/v) TFE, 20 mM sodium acetate, 150 mM NaCl (pH 4.5) solution with the same peptide and DSS concentrations. All direct dimension data was acquired using double quadrature detection. The ^15^N-Heteronuclear Single Quantum Correlation (HSQC) spectrum was collected with 1024 scans at a resolution of 2048 x 128 (TD_2_ x TD_1_) complex points in the direct (^1^H) and indirect dimension respectively over sweep widths of 7143 x 2431 Hz in the direct and indirect dimensions respectively. The acquisition mode for the indirect dimension was non-uniformly sampled Echo-Antiecho with fifty percent sampling and a 1 s recycling delay. The ^13^C-HSQC spectrum was collected with 64 scans at a resolution of 2048 x 256 complex points with sweep widths of 7143 x 9050 Hz. The acquisition mode for the indirect dimension was non-uniformly samples Echo-Antiecho with a fifty percent sampling and a 1 s recycling delay. The [^1^H, ^1^H]-TOCSY spectrum was collected with 64 scans at a resolution of 2048 x 256 complex points over a sweep width of 7143 x 7198 Hz. The acquisition mode for the indirect dimension was States-TPPI with a recycling delay of 1 s. Homonuclear Hartman-Hahn transfer was accomplished through an 80 ms MLEV17 mixing sequence. [^1^H, ^1^H]-Nuclear Overhauser Effect SpectroscopY (NOESY) spectra were collected with 48 scans at a resolution of 2048x480 complex points with sweep widths of 7143x5998Hz. States-TPPI acquisition was used with 48 scans and a 1 s recycling delay with water suppression achieved through a Watergate flip-back sequence. Spectra were acquired with mixing times of 100, 250, 400, and 500 ms. [^1^H, ^1^H]-Rotational Overhauser Effect SpectroscopY (ROESY) spectra were collected with 32 scans at a resolution of 2048 x 480 complex points with sweep widths of 7143 x 5998Hz. The acquisition mode for the indirect dimension was States-TPPI with a recycling delay of 1 s. The peptide was assigned in PBS and 90% TFE using a suite of 2D natural abundance experiments consisting of ^15^N- and ^13^C-HSQCs, TOCSYs, NOESYs, and ROESYs, all collected at 25C

### NMR Secondary Chemical Shift Calculation

Random coil chemical shift values for each residue type were obtained from BMRB^54^ tabulated values, with an average of 70k shift values for each amide proton, excluding proline.

### NMR Temperature Shift Calculation

1D ^1^H spectra were acquired at 10, 15, 20, 25, and 30C, each with 128 scans and a resolution of 12,288 points with a sweep width of 7143 Hz and a 2 s recycling delay. Chemical shift values for each amide resonance were fit as linear function of temperature in proFit.

## Supporting Information

Contains Table S1, Table S2, and Figure S1.

## AUTHOR CONTRIBUTIONS

All authors designed and performed the research, analyzed the data, and wrote the paper.

## ACKNOWLEDGMENTS

We would like to thank Dr. Hongwei Wu for his assistance collecting NMR data at the MetaCyt Biomolecular NMR Laboratory at Indiana University. We would also like to thank the staff at the Physical Biochemistry Instrumentation Facility (PBIF) at Indiana University as well for their assistance with collecting CD measurements. This work was supported by funding from the National Institutes of Health (R35GM143054, J.J.Z; T32DA024628, T.J.S.; T32GM140995, T.J.S.; and F31DA060484, T.J.S.) and German Research Foundation (CRC1423, Project 421152132, I.L.). MetaCyt and PBIF are supported by Indiana University and the State of Indiana.

