## Supplementary Information for "The isolated *Stachel* peptide of the adhesion G protein-coupled receptor GPR126 is intrinsically disordered"

| Gene | Stachel Peptide Sequence |
|---|---|
| ADGR-A1 | $^{1}$MDLKTVSLPRYPGE$^{15}$ |
| ADGR-A2 | $^{747}$GNVAVLMELSAFPREVGGAGAG$^{768}$ |
| ADGR-A3 | $^{738}$SNYAVLMDLTGELYTQAA$^{756}$ |
| ADGR-B1 | $^{927}$STFAILAQLSADADMEKATLPS$^{948}$ |
| ADGR-B2 | $^{912}$STFAVLAQPPKSLTLELAGSP$^{932}$ |
| ADGR-B3 | $^{857}$STFAILAQQPRELLMESSGTP$^{877}$ |
| ADGR-C1 | $^{2449}$ASFAVLMDISRRENGEVLPLKIVTY$^{2473}$ |
| ADGR-C2 | $^{2357}$TSFAVLMVSRRENGE$^{2372}$ |
| ADGR-C3 | $^{2518}$GTFGVLMDASPRERL$^{2532}$ |
| ADGR-D1 | $^{545}$TNFAILMQVVPLELA$^{559}$ |
| ADGR-D2 | $^{636}$TSFAILLQIYEVQRG$^{651}$ |
| ADGR-E1 | $^{585}$ANLAVIMASGELTMD$^{599}$ |
| ADGR-E2 | $^{518}$SSFAVLMAHYDVQEED$^{533}$ |
| ADGR-E3 | $^{339}$SSFAVLMALTSQEEDPVLTVITYV$^{362}$ |
| ADGR-E4P | $^{174}$SSFAVLVALAPKED$^{188}$ |
| ADGR-E5 | $^{531}$SSFAILMAHYDVEDYKLTLITR$^{552}$ |
| ADGR-F1 | $^{567}$TSFSILMSPFVPSTIFPVVKWITY$^{590}$ |
| ADGR-F2 | $^{430}$TSFSILMSPHILESLILTYIT$^{450}$ |
| ADGR-F3 | $^{753}$TAFSVLMSPHTVPEEPALALLTQ$^{775}$ |
| ADGR-F4 | $^{382}$SVVMSFSILMSSKS$^{395}$ |
| ADGR-F5 | $^{991}$TSFSILMSPDSPDPSSLLGILLD$^{1012}$ |
| ADGR-G1 | $^{383}$TYFAVLMVSSVEVDAVHLHYLSLLS$^{407}$ |
| ADGR-G2 | $^{277}$TSFAEPPDYSPVTHNV$^{292}$ |
| ADGR-G3 | $^{250}$TFFALLLRPTL$^{260}$ |
| ADGR-G4 | $^{2722}$THFGVLMDLSRSTV$^{2735}$ |
| ADGR-G5 | $^{227}$TYFAVLMQLSPALVPAELLAPLT |
| ADGR-G6 | $^{841}$THFGVLMDLPRSASQL$^{856}$ |
| ADGR-G7 | $^{416}$TNFAVLMTFRKDYQYPKSLDILSN$^{439}$ |
| ADGR-L1 | $^{839}$TNFAVLMAHRELYQGRINELLLS$^{861}$ |
| ADGR-L2 | $^{825}$TNFALMAHREIAYKD$^{840}$ |
| ADGR-L3 | $^{842}$TNFAVLMAHVEVKHSD$^{857}$ |
| ADGR-L4 | $^{407}$THFAILMSSGPSIG$^{420}$ |
| ADGR-V1 | $^{5891}$SVYAVYARTDNLSSYNE$^{5907}$ |

**Table S1.** *Stachel* peptide sequences for all 33 human aGPCRs. The *Stachel* peptide sequence of ADGR-A1 and ADGR-E4P are uncleavable**.**

| Gene | Protein Sequence |
| --- | --- |
| hADGR-G6 NTF-Stachel (aa 1-856) | MMFRSDRMWSCHWKWKPSPLLFLFALYIMCVPHSVWGCANCRVVLSNPSGTFTSPCYPNDYPNSQACMWTLRAPTGYIIQITF<br>NDFDIEEAPNCIYDSLSLDNGESQTKFCGATAKGLSFNSSANEMHVSFSSDFSIQKKGFNASYIRVAVSLRNQKVILPQTSDAY<br>QVSVAKSISIPELSAFTLCFEATKVGHEDSDWTAFSYSNASFTQLLSFGKAKSGYFLSISDSKCLLNNALPVKEKEDIFAESFE<br>QLCLVWNNSLGSIGVNFKRNYETVPCDSTISKVIPGNGKLLLGSNQNEIVSLKGDIYNFRLWNFTMNAKILSNLSCNVKGNVVD<br>WQNDFWNIPNLALKAESNLSCGSYLIPLPAAELASCADLGTLCQATVNSPSTTPPTVTTNMPVTNRIDKQRNDGIIYRISVVIQ<br>NILRHPEVKVQSKVAEWLNSTFQNWNYTVYVVNISFHLSAGEDKIKVKRSLEDEPRLVLWALLVYNATNNTNLEGKIIQ<br>QKLLKNNESLDEGLRLHTVNVRQLGHCLAMEEPKGYYWPSIQPSEYVLPCPDKPGFSASRICFYNATNPLVTYWGPVDISNCLK<br>EANEVANQILNLTADGQNLTSANITNIVEQVKRIVNKEENIDITLGSTLMNIFSNILSSSDSDLLESSSEALKTID<br>ELAFKIDLNSTSHVNITTRNLALSVSSLLPGTNAISNFSIGLPSNNESYFQMDFESGQVDPLASVILPPNLLENLSPEDSVLVRRA<br>QFTFFNKTGLFQDVGPQRKTLVSYVMACSIGNITIQNLKDPVQIKIKHTRTQEVHHPICAFWDLNKNKSFGGWNTSGCVAHR<br>DSDASETVCLCNHFTHFGVLMDLPRSASQL |
| hADGR-G6 GAIN-CTF (aa 665-1221) | DLNSTSHVNITTRNLALSVSSLLPGTNAISNFSIGLPSNNESYFQMDFESGQVDPL<br>ASVILPPNLLENLSPEDSVLVRRAQFTFFNKTGLFQDVGPQRKTLVSYVMACSIGNITIQ<br>NLKDPVQIKIKHTRTQEVHHPICAFWDLNKNKSFGGWNTSGCVAHRDSDASETVCLCNHF<br>THFGVLMDLPRSASQLDARNTKVLTFISYIGCGISAIFSAATLLTYVAFEKLRRDYPSKI<br>LMNLSTALLFLNLLFLLDGWITSFNVDGLCIAVAVLLHFFLLATFTWMGLEAIHMYIALV<br>KVFNTYIRRYILKFCIIGWGLPALVVSVVLASRNNNEVYGKESYGKEKGDEFCWIQDPVI<br>FYVTCAGYFGVMFFLNIAMFIVVMVQICGRNGKRSNRTLREEVLRNLRSVVSLTFLLGMT<br>WGFAFFAWGPLNIPFMYLFSIFNSLQGLFIFIFHCAMKENVQKQWRQHLCCGRFRLADNS<br>DWSKTATNIIKKSSDNLGKSLSSSSIGSNSTYLTSKSKSSSTTYFKRNSHTDNVSYEHSF<br>NKSGSLRQCFHGQVLVKTGPC |
| hADGR-G6 GAIN-bound (aa 665-856) | DLNSTSHVNITTRNLALSVSSLLPGTNAISNFSIGLPSNNESYFQMDFESGQVDPL<br>ASVILPPNLLENLSPEDSVLVRRAQFTFFNKTGLFQDVGPQRKTLVSYVMACSIGNITIQ<br>NLKDPVQIKIKHTRTQEVHHPICAFWDLNKNKSFGGWNTSGCVAHRDSDASETVCLCNHF<br>THFGVLMDLPRSASQL |
| hADGR-G6 GAIN-bound (aa 665-856) | THFGVLMDLPRSASQL |
| hADGR-G6 GAIN-CTF (aa 665-1221) | THFGVLMDLPRSASQLDARNTKVLTFISYIGCGISAIFSAATLLTYVAFEKLRRDYPSKI<br>LMNLSTALLFLNLLFLLDGWITSFNVDGLCIAVAVLLHFFLLATFTWMGLEAIHMYIALV<br>KVFNTYIRRYILKFCIIGWGLPALVVSVVLASRNNNEVYGKESYGKEKGDEFCWIQDPVI<br>FYVTCAGYFGVMFFLNIAMFIVVMVQICGRNGKRSNRTLREEVLRNLRSVVSLTFLLGMT<br>WGFAFFAWGPLNIPFMYLFSIFNSLQGLFIFIFHCAMKENVQKQWRQHLCCGRFRLADNS<br>DWSKTATNIIKKSSDNLGKSLSSSSIGSNSTYLTSKSKSSSTTYFKRNSHTDNVSYEHSF<br>NKSGSLRQCFHGQVLVKTGPC |

**Table S2.** Alphafold3 (upper two) and AIUPred (lower three) model constructs for h-ADGR-G6.

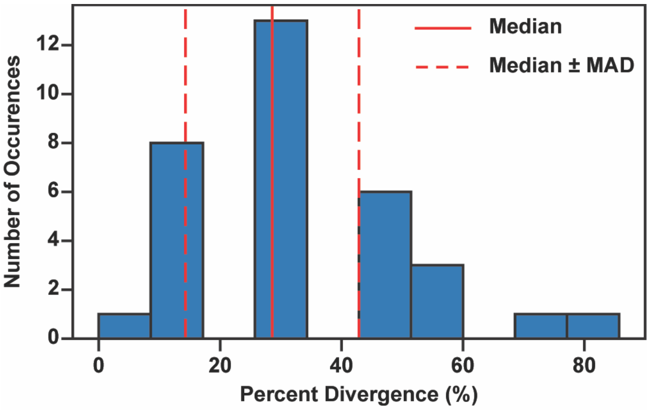


**Figure S1.** Plot of percent divergence (1-conservation score) for all 33 human aGPCR *Stachel* sequences defined as GPS+1 through TM1, residue 1.
